# Measuring the importance of annotation granularity to the detection of semantic similarity between phenotype profiles

**DOI:** 10.1101/086306

**Authors:** Prashanti Manda, James P. Balhoff, Todd J. Vision

## Abstract

In phenotype annotations curated from the biological and medical literature, considerable human effort must be invested to select ontological classes that capture the expressivity of the original natural language descriptions, and finer annotation granularity can also entail higher computational costs for particular reasoning tasks. Do coarse annotations suffice for certain applications? Here, we measure how annotation granularity affects the statistical behavior of semantic similarity metrics. We use a randomized dataset of phenotype profiles drawn from 57,051 taxon-phenotype annotations in the Phenoscape Knowledgebase. We compared query profiles having variable proportions of matching phenotypes to subject database profiles using both pairwise and groupwise Jaccard (edge-based) and Resnik (node-based) semantic similarity metrics, and compared statistical performance for three different levels of annotation granularity: entities alone, entities plus attributes, and entities plus qualities (with implicit attributes). All four metrics examined showed more extreme values than expected by chance when approximately half the annotations matched between the query and subject profiles, with a more sudden decline for pairwise statistics and a more gradual one for the groupwise statistics. Annotation granularity had a negligible effect on the position of the threshold at which matches could be discriminated from noise. These results suggest that coarse annotations of phenotypes, at the level of entities with or without attributes, may be sufficient to identify phenotype profiles with statistically significant semantic similarity.

## INTRODUCTION

To make phenotype descriptions in the biological and medical literature amenable to large-scale discovery and compuation, a variety of efforts have been launched to convert such descriptions into logical expressions using ontologies and to integrate them into the larger ecosystem of online, open biological information resources [1]. Typically, this involves curation and annotation of phenotypes in the Entity-Quality (EQ) formalism [2], which is widely used by model organism communities for representation of gene phenotypes [3]. The EQ formalism has more recently been adopted by the Phenoscape project to curate phenotypes from the literature that are reported to vary among evolutionary lineages [4] with the goal of linking them to gene phenotypes and generating hypotheses about the genetic bases of evolutionary transitions [5].

In the EQ approach, an entity represents a biological object, e.g. an anatomical structure, an anatomical space, or a biological process, while a quality represents a trait or property that an entity possesses, e.g, shape, color, or size. Curators often create complex logical expressions called post-compositions by combining ontology terms, relations, and spatial properties from multiple ontologies in different ways to create entities and qualities that adequately represent phenotypic descriptions. For example, “big supraorbital bone” is represented as E: *supraorbital bone* (UBERON_0004747), Q: *enlarged size* (PATO·0000586). A more complex description such as “parietal fused with supraoccipital bone” is represented by relating the two affected entities, *supraorbital bone* (UBERON_0004747) and *parietal* (UBERON_2001997) using the quality *fused with* (PATO_0000642).

Annotation of phenotypes at this level of ontological detail is time consuming and expensive [6]. Annotating evolutionary phenotypes at the finest level of granularity often requires curators to create new ontology terms and request those terms to be added to the ontology. Coarse annotation removes the need for ontology development by limiting curators to a small set of attribute level qualities already present in the ontology. Reducing the effort on curatorial tasks such as ontology development and data preparation improves the annotation rate from two characters per hour to 14 characters per hour [6]. Thus coarse annotation can be part of an efficient annotation workflow, and permit larger datasets to be curated for equivalent resources. In addition, reasoning over the combinatorial entity and quality ontology space for EQ annotations poses a serious computational challenge.

Given these competing considerations, what level of annotation granularity is optimal? The answer may depend on the particular application. For Phenoscape, a major goal is to be able to find sets of phenotypes that show greater semantic similarity than would be expected by chance when comparing sets of phenotypes from different biological domains (e.g. those observed in evolutionary lineages versus those induced by genetic manipulations in the laboratory) [5]. When comparing phenotypes with such different biological origins, we would not expect to see congruence in fine detail for a variety of reasons. For instance, even if the same or homologous genes have contributed to the two profiles, independent changes to those genes may underpin the phenotypes, they may be in lineages for which the genetic networks have diverged, and there may have been considerable evolutionary modification of the phenotype since its first origin. Even if two biological phenotypes are identical, the way in which the phenotypes are observed and described by independent researchers may lead to natural language descriptions, and thus profiles of annotations, that are quite different. With such weak matches, do finer annotations enable similarities to be detected, or are finer annotations superfluous or even distracting?

To explore this issue, we have conducted experiments to test the statistical sensitivity of semantic similarity at varying annotation granularity. Our approach involves simulating phenotype profiles by sampling from real annotations drawn from the Phenoscape Knowledgebase [5]. We measured similarity between profiles that shared all, some or none of their annotations, with the remainder drawn randomly from the population of annotations. We assessed the decline of semantic similarity to the point at which it could no longer be discriminated from random chance. This was done for four different semantic similarity statistics, and for three levels of annotation granularity.

## METHODS

### A. Semantic similarity metrics

The four semantic similarity statistics we have chosen represent extremes along two different dimensions by which semantic similarity metrics vary [7–10]. Edge-based semantic similarity metrics use the distance between terms in the ontology as a measure of similarity. Node-based measures use the Information Content of the annotations to the terms being compared and/or their least common subsumer. The similarity metrics we have chosen are based on Jaccard (edge-based) and Resnik (node-based) similarity, which are popular in biological applications (e.g. [11]). For each, we have one version that summarizes the distribution of pairwise similarities between two sets of annotations, and another that calculates a groupwise score directly.

1) *Jaccard similarity:* The Jaccard similarity (*sJ*) of two classes (*A, B*) in an ontology is defined as the ratio of the number of classes in the intersection of their subsumers over the number of classes in their union of their subsumers [12].

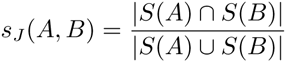

where *S*(*A*) is the set of classes that subsume A.

2) *Resnik similarity:* The Information Content of ontology class *A*, denoted *I*(*A*) is defined as the negative logarithm of the proportion of profiles annotated to that class *f* (*A*) out of *T* profiles in total.

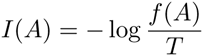

Since the minimum value of *I*(*A*) is zero, at the root of the ontology, while the maximum value is − log(1*/T*), we can compute a Normalized Information Content (*I*_*n*_) with range [0, 1]

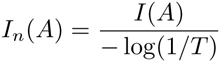

The Resnik similarity (*s*_*R*_) of two ontology classes is defined as the Normalized Information Content of the least common subsumer (LCS) of the two classes.

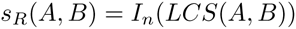

### B. Profile similarity

A set of ontology-based phenotype annotations is called a phenotype profile. When comparing two profiles, *X* and *Y*, where each has at least one, and potentially many annotations, we could either summarize all the pairwise combinations of annotations, or we could compute a groupwise similarity measure directly as a function of graph overlap.

*1) Best Pairs:* Pairwise approaches summarize the distribution of pairwise Jaccard or Resnik similarity scores between annotations in the two profiles. Here we use the Best Pairs score. For each annotation in *X*, the best scoring match in *Y* is determined, and the median of the |*X*| resultant values is taken. Similarly, for each annotation in *Y*, the best scoring match in *X* is determined, and the median of the |*Y*| values is taken. The Best Pairs score *p*_*z*_ (*X, Y*) is the mean of these two medians. The index *z* can be used to denote whether the pairwise values are Resnik (*z* = *R*) or Jaccard (*z* = *J*).

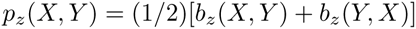

where

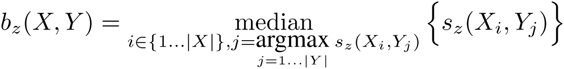

Note that, as defined, *p*_z_(*X, Y*) = *p*_z_ (*Y, X*).

*2) Groupwise:* Groupwise approaches compare profiles directly based on set operations or graph overlap.

The Groupwise Jaccard similarity of profiles *X* and *Y, g*_*J*_ (*X, Y*), is defined as the ratio of the number of classes in the intersection to the number of classes in the union of the two profiles

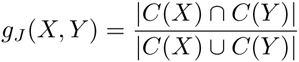

where *C*(*X*) is the set of classes belonging to *X* plus their subsumers.

Similarly, the Groupwise Resnik similarity of profiles *X* and *Y*, _*gR*_(*X, Y*), is defined as the ratio of the normalized information content summed over all nodes in the intersection of *X, Y* to the information content summed over all nodes in the union.

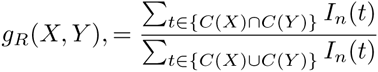

where *C*(*X*) is defined as above.

**Fig. 1.**
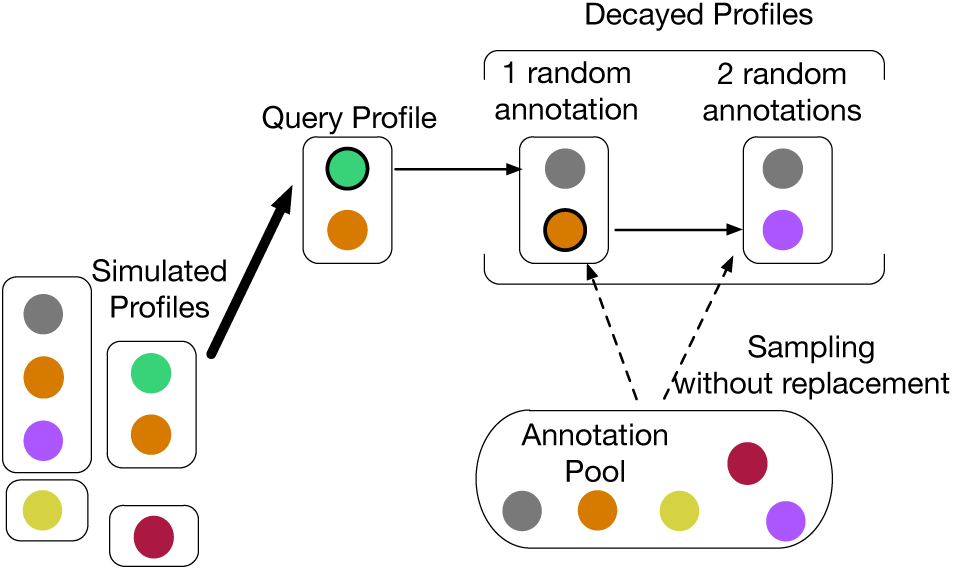
Profile decay via iterative replacement. Query profiles are selected from the pool of simulated profiles (lower left). Filled circles represent annotations, and annotations within the same profile are enclosed by boxes. Circles of the same color represent the same annotation. At each iteration, one of the remaining original annotations in the query profile is replaced with a randomly selected annotation from the pool. The process continues until each of the annotations in the original query profile has been replaced.

### C. Source data

The Phenoscape Knowledgebase contains a dataset of 661 taxa with 57,051 evolutionary phenotypes, which are phenotypes that have been inferred to vary among the taxon’s immediate descendents [5]. A simulation dataset of subject profiles having the same size distribution of annotations per taxon was created by permutation of the taxon labels.

### D. Simulating profile ‘decay’

To simulate decay of profile similarity, five query profiles of size ten were randomly selected from the simulated dataset. For each, there is one profile among the set of subjects for which each annotation has a one-to-one perfect match. For each of the five profiles, ten progressively decayed profiles were obtained by iteratively replacing one of the original annotations with an annotation randomly selected from among the 57,051 available (Figure II-D). Thus, for each original profile, there is a profile in which one original annotation has been replaced with random annotation, another in which two have been replaced, and so on, through to a fully decayed profile in which all original annotations have been replaced with a random one. To characterize the noise distribution for each metric in the absence of semantic similarity, we also generated 5,000 profiles of size ten by drawing annotations randomly from among the 57,051 available. These profiles would not be expected to have more than nominal similarity with any of the simulated subject profiles.

### E. Adjusting annotation granularity

The evolutionary phenotypes available from Phenoscape have been annotated with both entities and qualities, and the intermediate level of attribute is implicit in the quality annotation due the structure of the PATO quality ontology [4]. In order to measure semantic similarity for three levels of granularity: entity only (E), entity-attribute (EA), and entity-quality (EQ), we used three different phenotype ontologies, one for each granularity level, containing phenotype concepts combining terms from Uberon (entities) and PATO (attributes and qualities). In each evaluation, annotations in the query profiles and the simulated database will match at the granularity level available in the generated phenotype ontology.

## RESULTS AND DISCUSSION

We measured semantic similarity between each of the five query profiles and their decay series to all 661 profiles in the subject database. This was done for each of the four semantic similarity metrics (Best Pairs and Groupwise variants of Jaccard and Resnik metrics) and for each of the three granularity levels (E: Entity only, EA: Entity-Attribute, and EQ: Entity-Quality). The results are shown in Figure 2). For ease of interpretation, we take the upper 99.9% of the similarity distribution for random profile matches as an arbitrary threshold for comparing the sensitivity of the different series. All series cross this threshold when approximately half of the annotations have been replaced, with a sudden decline in similarity for the Best Pairs statistics and a more gradual decline for the groupwise statistics. While the differences in sensitivity among the annotation granularity levels are subtle, the annotations of intermediate granularity (EA) have marginally greater sensitivity for all four statistics.

**Fig. 2.**
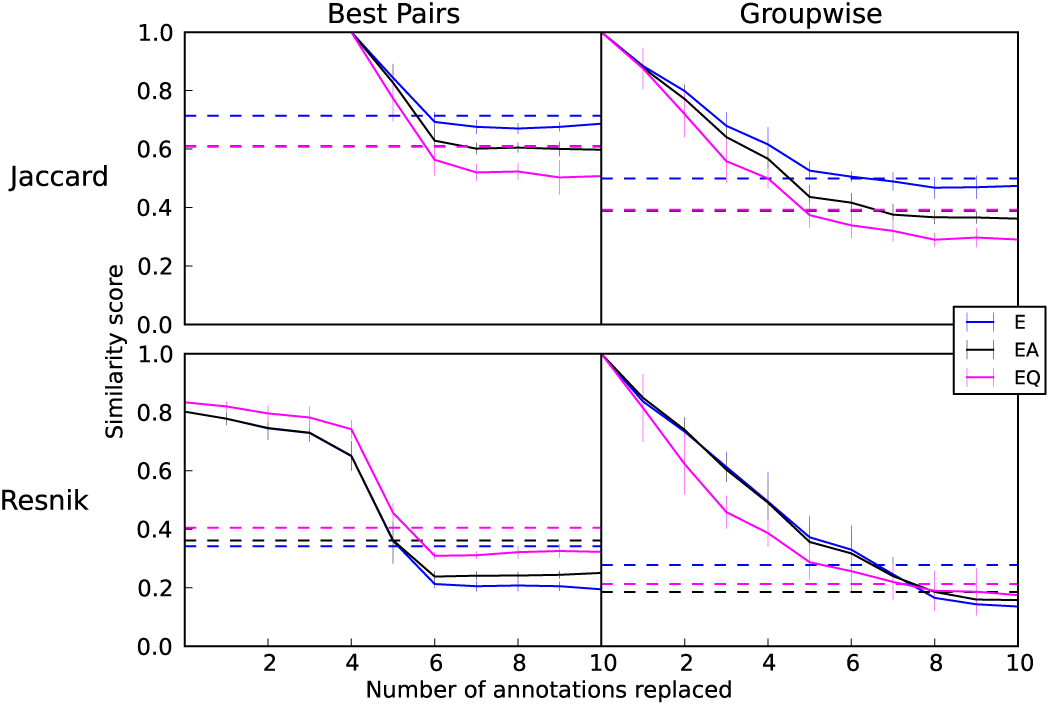
Pattern of similarity decay with E, EA, and EQ data as profiles are decayed via Random Replacement. Solid lines represent the mean best match similarity of the 5 query profiles to the database after each annotation replacement. Error bars show 2 standard errors of the mean. Dotted lines represent the 99.9th percentile of the noise distribution.

The sharp decline in similarity under the Best Pairs statistics at approximately 50% decay can be understood as a result of summarizing the pairwise distribution with the median. In future work, we aim to explore how the sensitivity of pairwise statistics might be tuned by using different percentiles. Given the relatively flat performance of the Best Pairs statistics when decay was under 50%, we suggest that groupwise statistics are likely to provide greater discrimination between true matches of varying quality and thus better for rank ordering the outcome of semantic similarity searches, e.g. [5]. Our results also illustrate how difficult it can be to statistically discriminate weakly matching profiles from noise, something which has received relatively little consideration in many applications of semantic similarity search to date.

The relatively minor differences in statistical performance with varying annotation granularity, with EA showing marginally greater sensitivity, has implications both for the process of generating annotations and the implementation of semantic similarity computation. As noted in the Introduction, annotation to EA requires considerably less human curation effort than EQ, and is almost identical in effort to curation to E. Restricting annotation granularity to EA may also ease the challenge of speeding of curation through machine-aided natural language processing, e.g. [13].

Second, the computational expense of measuring semantic similarity can be prohibitive for fine-grained annotations due to an explosion in the number of classes required for reasoning when annotations draw from multiple ontologies [14]. If the inclusion of qualities does not improve sensitivity, that opens up the possibility of conducting fast, scalable on-the-fly web-based semantic searches at coarser annotation levels.

One of the contributions of this work is in introducing a framework for evaluating the statistical sensitivity of semantic similarity metrics. Nonetheless, the results reported here are specific to one particular model for the decay of similarity between two profiles, in which some portion of annotations that match perfectly while others do not match at all. We recognize a need to explore other models, especially ones where pairs of annotations may match imperfectly. We also propose that other evaluation criteria should be examined to more fully understand the trade-offs involved in building datasets with a particular level of annotation granularity.

## ACKNOWLEDGEMENTS

We thank W. Dahdul, T.A. Dececchi, N. Ibrahim and L. Jackson for curation of the original dataset, along with the larger community of ontology contributors and data providers (http://phenoscape.org/wiki/Acknowledgments#Contributors), and useful feedback from P. Mabee, H. Lapp, W. Dahdul, and other members of the Phenoscape team. This work was funded by the National Science Foundation (DBI-1062542).

